# uniSynK: a ratio-optimized bicistronic chemically induced dimerization tool for robust induction of dendritic spine enlargement

**DOI:** 10.64898/2026.05.02.720486

**Authors:** Kota Homma, Masanari Ohtsuka, Siqi Zhou, Hitoshi Okazaki, Tomoki Arima, Haruo Kasai, Takeshi Sawada

## Abstract

Synaptic plasticity is widely implicated in learning and brain-state regulation. Although correlations between synaptic modifications and diverse brain functions have been extensively reported, establishing causality requires tools that can directly and selectively perturb synapses in vivo. Recently, the chemogenetic actuator SYNCit-K (SynK) enabled causal induction of dendritic spine enlargement and synaptic potentiation. However, SynK relies on the co-expression of two independent components delivered by separate viral vectors, making it difficult to reliably achieve appropriate expression levels of both components within the same neuron. Here, using mouse neocortical dissociated culture, we sought to develop a unified, single-vector version of SynK (uniSynK) that enables robust and safe induction of dendritic spine enlargement. We first found that each SynK component induces distinct, concentration-dependent abnormalities in neuronal morphology when overexpressed, underscoring the need for precise control of their expression ratio. To address this challenge, we engineered a single construct in which the two components are linked by internal ribosome entry site (IRES) variants with graded translational efficiencies. Systematic screening of IRES variants identified an optimized configuration that preferentially drives each component within its respective concentration range that preserves normal spine morphology. The optimized uniSynK efficiently induces dendritic spine enlargement, while exhibiting fewer off-target morphological abnormalities than observed in the multi-vector system when expression levels were not properly controlled. By simplifying experimental design, this single-vector, ratio-optimized synaptic chemogenetic tool provides a practical and broadly applicable platform that should facilitate causal interrogation of synaptic plasticity across neural circuits and behavior.

**Significance Statement:** Synaptic plasticity is thought to shape learning, brain states, and neuropsychiatric disease, but testing its causal roles requires tools that can directly manipulate synapses with minimal side effects. Here, we developed uniSynK, a single-vector chemogenetic system that induces dendritic spine enlargement by coordinating the expression of two functional components. By defining safe expression ranges and using IRES variants to optimize their ratio, uniSynK achieves robust spine enlargement while reducing unintended morphological abnormalities associated with imbalanced expression. This work provides both a practical tool for synaptic manipulation and a general strategy for optimizing multi-component biological systems.

## Introduction

Synaptic plasticity is a fundamental mechanism through which the brain adapts to experience. Long-term potentiation (LTP), for example, has been intensively studied as a cellular correlate of learning (Kasai et al., 2021; Matsuzaki et al., 2004; Xu et al., 2009), whereas alterations in dendritic spines—the structural substrates of excitatory synapses—including changes in number, morphology, and associated genes, have been linked to psychiatric conditions (McCann and Ross, 2017; Penzes et al., 2011). Recent studies have further suggested that synaptic plasticity is dynamically regulated across brain states, with prolonged wakefulness leading to dendritic spine enlargement and accumulation of excitatory synaptic receptors (de Vivo et al., 2017; Diering et al., 2017).

Despite these associations, the causal roles of these synaptic changes remain incompletely understood. Addressing this challenge requires tools that enable selective manipulation of defined synapses. Optogenetic approaches have been widely developed over the past two decades, with a subset adapted to manipulate synaptic plasticity (Goto et al., 2021; Hayashi-Takagi et al., 2015; Murakoshi et al., 2017; Shibata et al., 2021). While offering high spatial and temporal resolution, they are less suited for chronic or large-scale manipulation due to limitations such as tissue heating and the need for invasive optical hardware.

To circumvent these limitations, a chemogenetic tool to induce dendritic spine enlargement and correlated synaptic potentiation (Holler et al., 2021; Matsuzaki et al., 2004), SYNCit-K (Synapse-targeted Chemically induced translocation of Kalirin-7; SynK) was recently developed (Sawada et al., 2024). SynK employs rapamycin-dependent FKBP–FRB dimerization (Inoue et al., 2005) to mimic the physiological shuttling of Kalirin-7, a Rac1-activating guanine nucleotide exchange factor (GEF), into postsynaptic densities (PSD) (Xie et al., 2007), thereby enlarging dendritic spines and strengthening excitatory synapses. Importantly, SynK enables causal interrogation of synaptic potentiation at the behavioral level, as demonstrated by increased NREM sleep and δ-power following SynK-induced potentiation of prefrontal cortical synapses (Sawada et al., 2024).

Despite these advances, the limited packaging capacity of AAV (less than 4.7 kb) (Halbert et al., 2002; Tornabene et al., 2019) makes it challenging to accommodate the two components —one encoding a truncated form of Kalirin-7 (Kal7ΔN) conjugated with FKBP (FKBP-Kal7ΔN) and the other a postsynaptic density (PSD)-anchored FRB array (PSDΔ1,2-FRB)— such that the current system relies on two separate AAVs.

AAV-mediated expression typically varies across infected neurons and across the infection site (Chen et al., 2013; Tian et al., 2009). As a result, achieving an appropriate expression level—sufficient to elicit an effect while avoiding overexpression-related side effects (Dana et al., 2014; Goossens et al., 2021; Mori et al., 2020; Moriya, 2015)—in a large proportion of cells remains challenging. This challenge is exacerbated when multiple AAVs are used, as coordinated expression of all components at appropriate levels is required within the same cells, thereby reducing the fraction of cells meeting these conditions and complicating the application of the system (Grieger et al., 2006; Maturana et al., 2023), a problem that may be further compounded by variability across AAV preparations. One strategy to mitigate this limitation is to place both components within a single AAV construct by compacting the design, thereby coupling their expression and facilitating coordinated control of their levels.

In this study, we first attempted to shorten the coding sequences of the two SynK components to enable packaging into AAV vectors, using dissociated neuronal cultures. We then determined the maximum expression levels of each component that did not cause off-target effects in baseline conditions. To further streamline the system, we redesigned SynK as a single-vector construct by linking the two components via an internal ribosome entry site (IRES). By selecting IRES variants with graded translational efficiencies (Koh et al., 2013), we fine-tuned the FKBP/FRB expression ratio, allowing each component to be expressed near its maximal level without detectable off-target effects. Notably, the optimized unified-SynK construct (uniSynK) robustly induced dendritic spine enlargement, while exhibiting fewer side effects than the original split-vector design which exhibited variability in the expression levels of the two components. This one-vector, ratio-optimized uniSynK should simplify future in vivo applications and provide a practical platform for causal interrogation of synapse–behavior relationships.

## Materials and Methods

All animal procedures followed the guidelines of the Animal Experimental Committee of the Faculty of Medicine at the University of Tokyo.

### Plasmid and AAV preparation and purification

Plasmids were constructed using standard molecular cloning techniques, including restriction enzyme digestion, PCR amplification, and In-Fusion assembly (Takara Bio). EMCV IRES variants (v10, v12) (Koh et al., 2013) were generated by PCR-based mutagenesis. All constructs were sequence-verified and prepared using endotoxin-free plasmid purification kits (Qiagen).

pCAG-mScarlet-C1

pAAV-CaMKII(0.3)-DIO-mScarlet-WPRE

pAAV-CaMKII(0.4)-Cre-WPRE-hGHpolyA

pAAV-CaMKII(0.3)-PSDΔ1,2-FRB-WPRE-hGHpolyA

pAAV-CaMKII(0.3)-PSDΔ1,2-FRB-mTq-WPRE-hGHpolyA

pAAV-CaMKII(0.3)-mVenus-FKBP-Kalirin7GEF-Cterm-WPRE-hGHpolyA

pAAV-CaMKII(0.3)-mVenus-FKBP-Kalirin7GEF-WPRE-hGHpolyA

pAAV-CaMKII(0.3)-PSDΔ1,2-FRB-mTq-IRESwt-mVenus-FKBP-Kalirin7GEF-W3SL

pAAV-CaMKII(0.3)-PSDΔ1,2-FRB-mTq-IRESv10-mVenus-FKBP-Kalirin7GEF-W3SL

pAAV-CaMKII(0.3)-PSDΔ1,2-FRB-mTq-IRESv10-mVenus-FKBP-Kalirin7GEF(K1393A)-W3SL

pAAV-CaMKII(0.3)-PSDΔ1,2-FRB-mTq-IRESv12-mVenus-FKBP-Kalirin7GEF-W3SL

pAAV-CaMKII(0.3)-PSDΔ1,2-FRB-IRESv10-FKBP-Kalirin7GEF-W3SL

AAVs were produced and their titers were measured as described previously (Grieger et al., 2006). In brief, plasmids for the AAV vector, pHelper (Stratagene) and pUCmini-iCAP30 PHP.eB (#103005, Addgene) were transfected into HEK293 cells (AAV293, Stratagene). After three days, cells were harvested and AAVs were purified twice using iodixanol. The titers of AAVs were estimated using quantitative PCR. uniSynK plasmid has been deposited at Addgene.

### Dissociated culture of neocortex

Dissociated cortical neuron cultures were prepared from embryonic day 17 (E17) ICR mouse embryos (SLC) of both sexes (Okabe et al., 1999). Cells were plated at a density of 4.0 × 10⁴ cells/cm² on poly-L-lysine-coated glass-bottom dishes (Greiner, Advanced TC). Cultures were maintained at 37 °C in a humidified atmosphere with 5% CO₂ in Neurobasal medium supplemented with B27 (1:50; Gibco) and L-glutamine (1:200; Gibco). Half of the medium was replaced every ∼5 days.

### Plasmid lipofection and AAV transduction to dissociated cortical neurons

For plasmid delivery via lipofection, plasmid mixtures were combined with Lipofectamine 2000 (Invitrogen) and incubated for 20 min. Neurons were transfected at DIV16, and the medium was replaced after 4 h with a 1:1 mixture of pre-collected conditioned medium and fresh culture medium. For AAV transduction, viral mixtures were added to neurons at DIV11-12, and the medium was replaced with a 1:1 mixture of conditioned medium and fresh culture medium, after 20-24h.

### Dendritic spine imaging in dissociated cultures and image processing

Time-lapse imaging of up to three fluorescent channels (mTurquoise2, mVenus, and mScarlet) was performed every 20 min using a confocal microscope (A1R, Nikon), equipped with a 60× oil-immersion objective lens (NA 1.40). Z-stack images were acquired with a step size of 0.3 μm along the z-axis, typically spanning ∼30 optical sections to fully encompass the analyzed spines. To activate SynK via induced dimerization, half of the culture medium was removed and retained in advance. Immediately before application, a portion of the remaining medium was mixed with the A/C heterodimerizer (Takara Bio; 0.5 mM stock in ethanol) and then returned to the dish to achieve a final concentration of 4 μM. After 10 min, cells were returned to the retained conditioned medium.

Representative images were rotated when necessary using bilinear interpolation in ImageJ. Spines with more than two spinules were classified as highly branched (Penzes et al., 2001), and their fraction was calculated per dendrite (Fig. 4F).

### Measurement of time series changes in spine volume

Time-lapse z-projected images were aligned using the ITK registration framework in Python. Signal leakage across channels was quantified using neurons expressing only mScarlet, mVenus, or mTurquoise2 (mTq), and these values were used for spectral unmixing. After processing, the background-subtracted mScarlet fluorescence intensity of each spine head, normalized to the total fluorescence of the imaging area, was used as a proxy for spine-head volume, as described previously (Matsuzaki et al., 2004). Spines (4–20 per dendrite) were randomly selected for analysis. Spines that could not be consistently enclosed within the same ROI across all time points were excluded. In addition, spines lacking detectable mTq signal were excluded from mVenus/mTq ratio analysis. Baseline spine volume was defined as the mean across all time points prior to A/C heterodimerizer application. For quantification and statistical analyses, relative increases in spine volume from 20 to 120 min after A/C heterodimerizer administration were averaged.

In experiments with non-sparse AAV-mediated expression of mVenus and mTq (Fig. 7), to minimize signal contamination from neighboring neurons, a mask was generated by applying 2D Otsu thresholding to the cell-fill mScarlet channel. Regions lacking signal were excluded from the other channels (mTq and mVenus) prior to alignment of z-projected images, followed by all subsequent processing described above.

### Measurement of absolute dendritic spine volume and its intrinsic fluctuations

To estimate the absolute spine volume, z-stack images were deconvolved in Python using flowdec with the Richardson–Lucy method based on a synthetic point spread function calculated from imaging parameters, and then binarized using Otsu’s method. The absolute spine volume was calculated by counting the voxels within the binarized spine regions and multiplying the voxel count by the voxel volume. For the analysis of intrinsic spine fluctuations (Fig. 3B-D and 6K, L), the mean change in spine volume (μ) and the standard deviation (σ) were calculated over 20-min intervals at 45–25 min and 25–5 min before, and 35–55 min after A/C heterodimerizer administration, as previously described (Ishii et al., 2018).

### Statistical analysis

Sample sizes were determined empirically based on previous studies and are comparable to those generally used in the field. Data are presented as individual data points with mean ± s.e.m., where applicable. Statistical analyses were performed using nonparametric methods, with Bonferroni correction applied where appropriate for multiple comparisons. P values are reported as unadjusted values, and statistical significance was assessed using corrected significance thresholds.

## Results

### Kalirin-7 C-terminal truncation and fluorescent protein addition do not impair SynK function

To rationally reduce the size of the SynK constructs while preserving its functional core, we revisited the molecular requirements underlying Kalirin-7–mediated spine enlargement. The original SynK contained the DH-PH (GEF module) domain and the C-terminal region containing the PDZ ligand of Kalirin-7 (Fig. 1A-C) (Sawada et al., 2024). While the GEF module is directly responsible for Rac1 activation (Xie et al., 2007), the primary role of the PDZ ligand has been proposed to be subcellular targeting (Lee and Zheng, 2010). Thus, we hypothesized that the PDZ ligand is dispensable in the SynK system, in which rapid and stable recruitment of Kalirin-7 to dendritic spines is artificially achieved via the rapamycin analog (A/C heterodimerizer). To directly test this hypothesis, we removed the C-terminal region from the original construct (Fig. 1D). As a result, the truncated SynK construct expressed in dissociated cortical neurons (PSDΔ1,2-FRB, 3 μg/mL; mVenus-FKBP-K7GEF, 0.05 μg/mL) showed K7GEF translocation and spine enlargement (Fig. 1E, F) that were indistinguishable from those induced by the original construct (Fig. 1H), thereby defining the simplified functional unit required for SynK–mediated spine enlargement. All subsequent development was based on this truncated construct.

**Figure 1.**
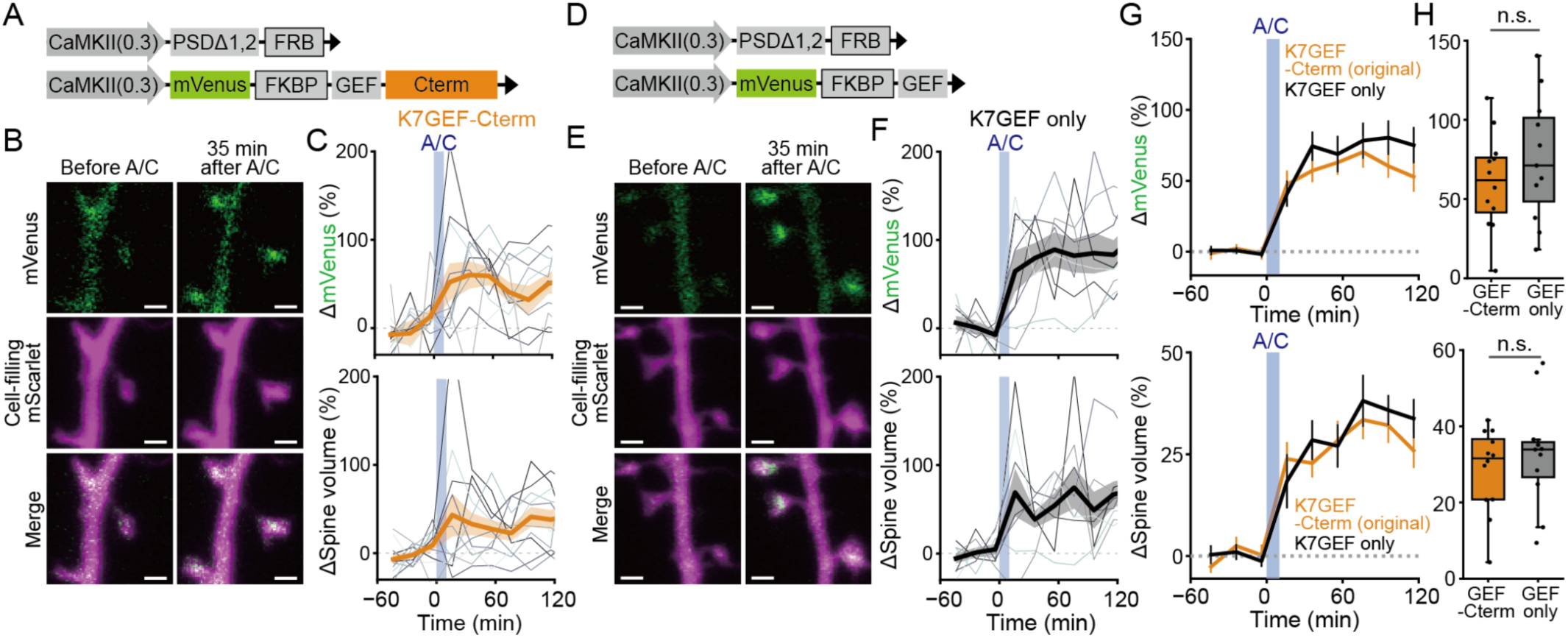
C-terminal truncation of Kalirin-7 preserves spine enlargement in SynK. ***A,*** Schematic of the original SynK construct. ***B,*** Representative confocal image of a dendrite expressing PSDΔ1,2–FRB (3 μg/mL) and mVenus–FKBP fused to Kalirin-7 GEF and C-terminal regions (0.05 μg/mL) before (−5 min) and after (35 min) application of A/C heterodimerizer. mScarlet signal is shown as a volume marker. Scale bar, 1 µm. ***C,*** Time courses of changes in the mVenus signal in dendritic spines (top) and spine volume (bottom), including their averages (bold line) and s.e.m (shade) for a representative neuron expressing PSDΔ1,2–FRB and mVenus–FKBP–K7GEF–Cterm. Values are plotted as a percentage change from the baseline average. ***D,*** Plasmid construct of SynK lacking the Kalirin-7 C-terminal region. ***E,*** Representative confocal image of a dendrite expressing PSDΔ1,2–FRB (3 μg/mL) and mVenus–FKBP–K7GEF (w/o Cterm, 0.05 μg/mL). Scale bar, 1 µm. ***F,*** Time course of mVenus and mScarlet signals in dendritic spines for a representative neuron expressing PSDΔ1,2–FRB and mVenus–FKBP–K7GEF. ***G, H,*** Averaged time courses (***G***) and summary (***H***) of changes in mVenus signal in dendritic spines (top) and spine volume (bottom) from neurons expressing SynK with either K7GEF–C-terminal region (original) or K7GEF alone. Data in (***G***) are shown as the mean ± s.e.m. Values in (***H***) represent the average between 35 and 115 min after A/C heterodimerizer administration. N (dendrites/neurons) = 12/7 (K7GEF-Cterm) and 11/4 (K7GEF only). The median (horizontal line), quartiles (boxes), and range within 1.5 times the interquartile range (whiskers) are denoted. Statistical comparisons were made using the Mann–Whitney test (top: *U* = 53, *p* = 0.44; bottom: *U* = 58, *p* = 0.64). \**p* < 0.05; \*\**p* < 0.01; n.s., not significant.

To quantitatively assess how the expression levels of the two components influence efficacy and potential side effects, we next modified the system so that both components were fluorescently labeled, fusing mTq to PSDΔ1,2-FRB, while the FKBP-K7GEF component retained its original mVenus tag (Fig. 2A). In neurons expressing PSDΔ1,2-FRB-mTq, the mTq signal was localized to dendritic spines (Fig. 2B). Following A/C heterodimerizer administration, FKBP-mVenus robustly translocated to dendritic spines, and spine enlargement was comparable to that observed with the original SynK constructs (Fig. 2B-E). These results indicate that mTq tagging does not interfere with PSDΔ1,2-FRB localization and preserves the functionality of the SynK system.

**Figure 2.**
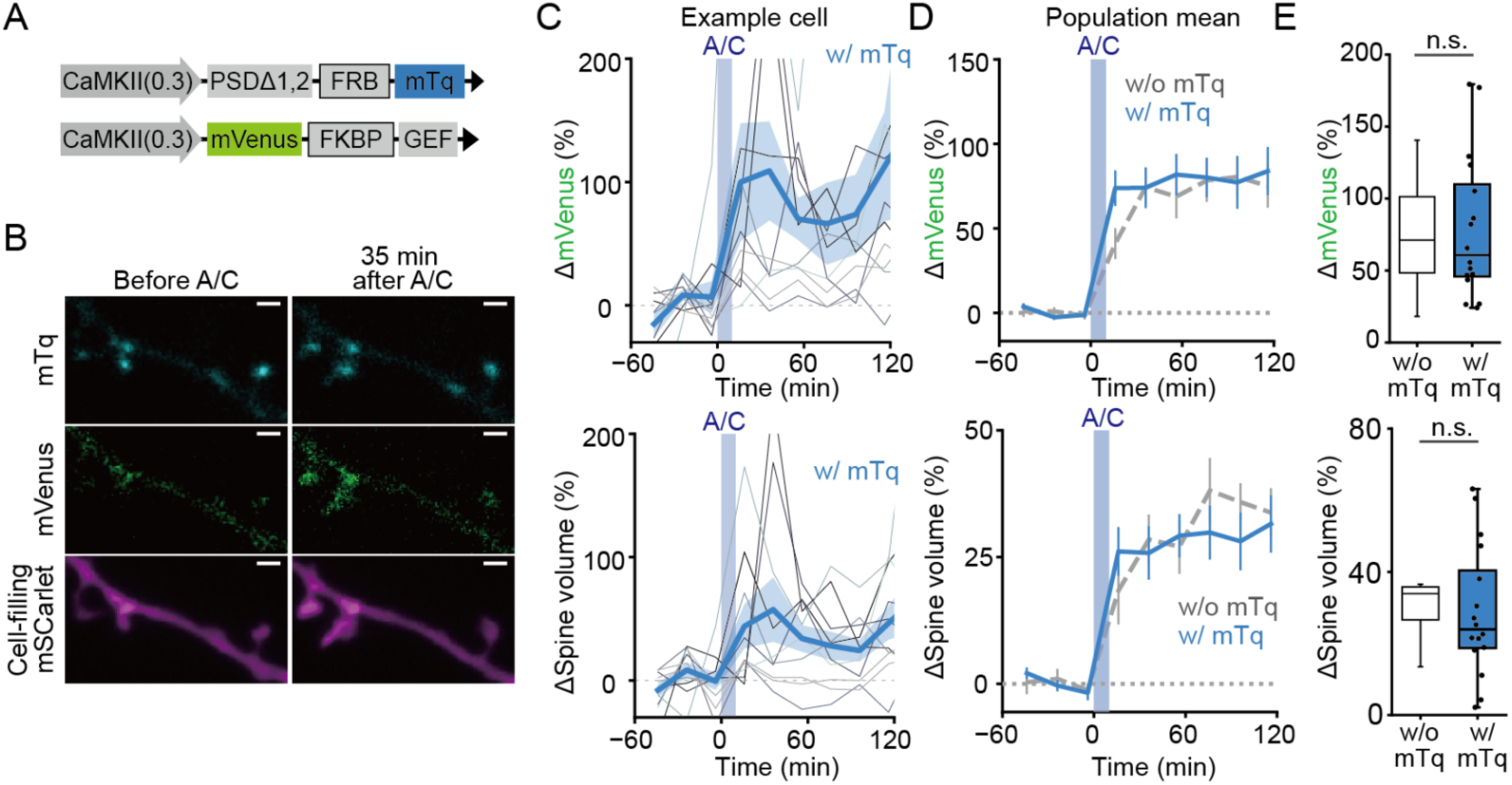
mTq tagging of PSDΔ1,2-FRB does not interfere with SynK function. ***A,*** Schematic of the SynK construct with mTq fused to PSDΔ1,2–FRB and lacking the Kalirin-7 C-terminal region. ***B,*** Representative confocal images of a dendrite expressing PSDΔ1,2-FRB-mTq and mVenus-FKBP-K7GEF before (−5 min) and after (35 min) application of A/C heterodimerizer. Images of mScarlet signal are shown as a volume marker. Scale bar, 1 µm. ***C***, Time courses of changes in the mVenus signal in dendritic spines (top) and spine volume (bottom), including their averages (bold line) and s.e.m (shade) for a representative neuron expressing PSDΔ1,2–FRB–mTq and mVenus–FKBP–K7GEF–Cterm. Values are plotted as a percentage change from the baseline average. ***D, E,*** Averaged time courses (***D***) and summary (***E***) of changes in mVenus signal in dendritic spines (top) and spine volume (bottom) from neurons expressing SynK with mTq tagging of PSDΔ1,2-FRB. Data without mTq are reproduced from Fig. 1 and shown as dashed lines (***D***) and hollow box plots (***E***) for comparison. Data in (***D***) are shown as the mean ± s.e.m. Values in (***E***) represent the average between 35 and 115 min after A/C heterodimerizer administration. N (dendrites/neurons) = 16/7 (w/ mTq) and 11/4 (w/o mTq). The median (horizontal line), quartiles (boxes), and range within 1.5 times the interquartile range (whiskers) are denoted. Statistical comparisons were made using the Mann–Whitney test (top: *U* = 91, *p* = 0.90; bottom: *U* = 106, *p* = 0.39). \**p* < 0.05; \*\**p* < 0.01; n.s., not significant.

Compared with the original construct (Sawada et al., 2024), removal of the C-terminal region and addition of mTq did not result in more pronounced spine hypertrophy prior to A/C administration (Fig. 3A), phenotypes typically observed upon Kalirin-7 overexpression (Xie et al., 2007). In addition, no significant changes in spine fluctuation (Ishii et al., 2018; Minerbi et al., 2009; Yasumatsu et al., 2008) were detected either before or after A/C administration in neurons expressing PSDΔ1,2-FRB-mTq and mVenus-FKBP-K7GEF (Fig. 3B-D).

**Figure 3.**
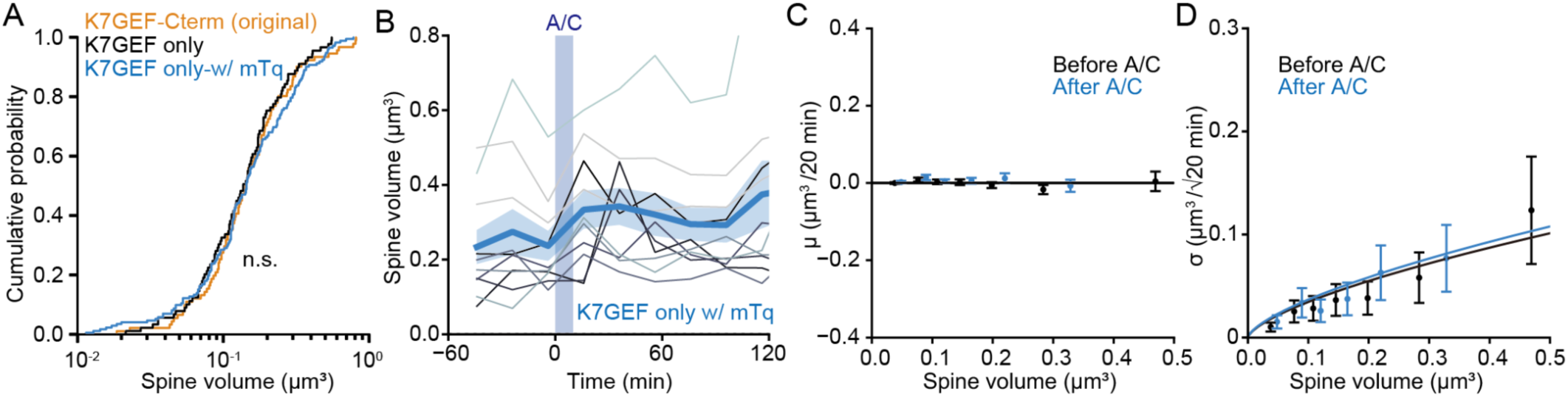
Removal of the Kalirin-7 C-terminal region and addition of mTq does not induce significant changes in spine volume distribution or intrinsic dynamics. ***A,*** Cumulative distribution of dendritic spine volumes under three conditions. K7GEF-Cterm (original), N = 92; K7GEF only, N = 89; K7GEF only-w/ mTq, N = 172 spines. Statistical comparisons were made using the Kolmogorov-Smirnov (K-S) test with Bonferroni correction. K7GEF-Cterm vs K7GEF only, *D* = 0.065, *p* = 0.97; K7GEF-Cterm vs K7GEF only-w/ mTq, *D* = 0.085, *p* = 0.74. ***B***, Time courses of the estimated absolute volume (see Methods) of spines before and after A/C heterodimerizer application, along with their averages (blue line) and s.e.m. (blue shade) for a representative neuron expressing PSDΔ1,2-FRB-mTq and mVenus-FKBP-K7GEF. ***C, D,*** Intrinsic spine fluctuation of neurons expressing PSDΔ1,2-FRB-mTq and mVenus-FKBP-K7GEF, before and after A/C heterodimerizer administration, each calculated per 20-min interval. Each plotted point represents the mean (***C***) and the s.d. (***D***) of spine-head volume changes in 24 pooled spines with similar baseline volumes. Error bars represent s.e.m. values (***C***) and the 95% confidence intervals of the estimated s.d. (***D***). Data are fitted by a zero (***C***) and to the 2/3 power of the baseline spine volume (***D***). \**p* < 0.05; \*\**p* < 0.01; n.s., not significant.

Collectively, these data indicate that neither truncation of the C-terminal region of Kalirin-7 nor the addition of fluorescent proteins significantly compromises SynK-mediated spine enlargement or introduces detectable morphological abnormalities compared with the original construct.

### The two SynK components each induce concentration-dependent unintended effects on spine morphology under baseline conditions

In developing a single-vector system that enables expression of both components while minimizing unintended side effects, we first sought to define the safe expression range for each component individually. Primary cortical neurons were transfected with either mVenus–FKBP–K7GEF (Fig. 4) or PSDΔ1,2–FRB–mTq (Fig. 5). To assess potential side effects across a broad range of expression levels and independently of method-specific influences, we employed both DNA lipofection and AAV-mediated gene delivery.

**Figure 4.**
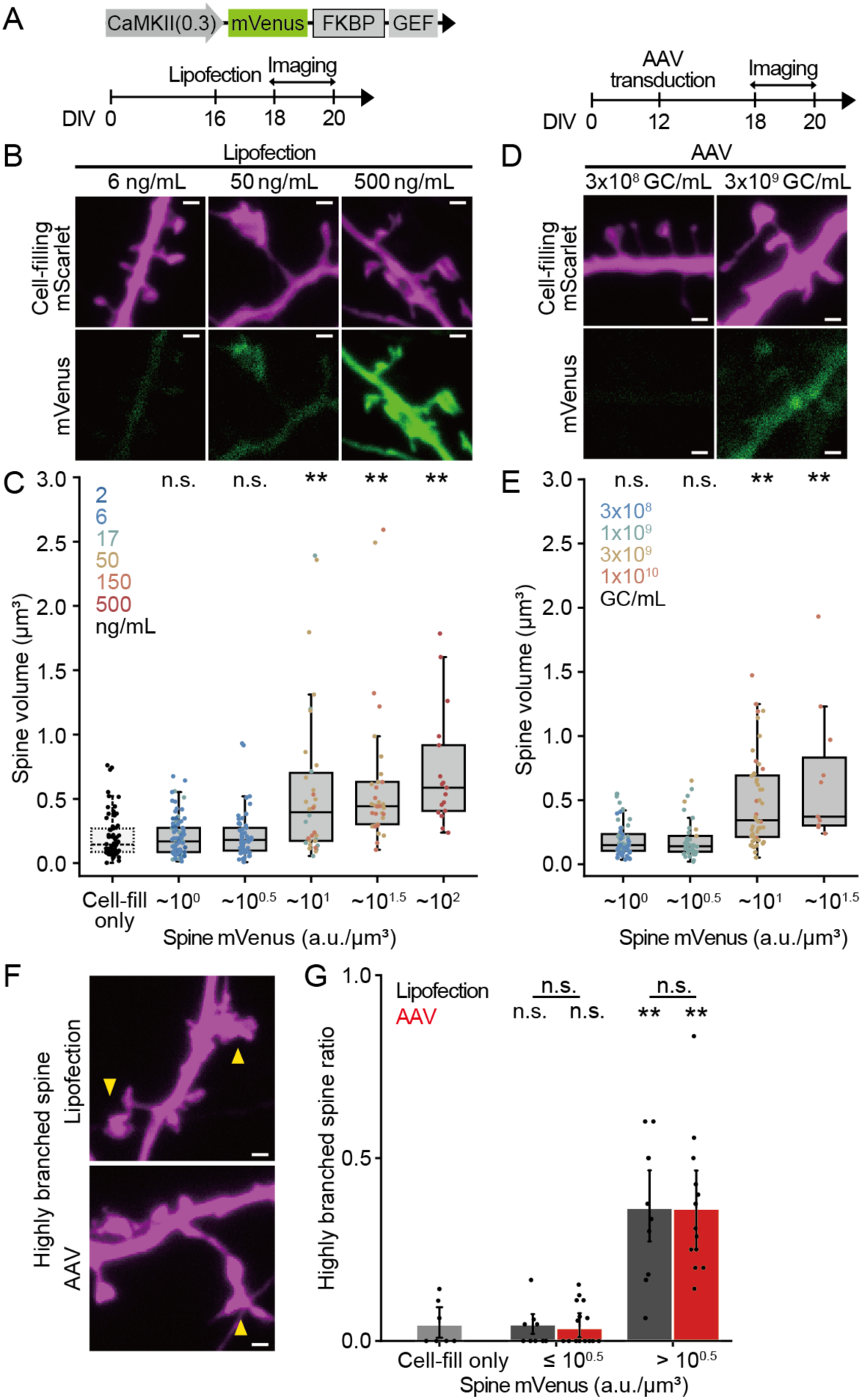
The Kalirin-7 GEF domain induces a concentration-dependent increase in baseline spine volume. ***A,*** Construct and experimental schedule. Two delivery methods were used: lipofection (***B, C***) and AAV (***D, E***). ***B,*** Example confocal images of dendrites expressing mVenus-FKBP-K7GEF via plasmid lipofection. mScarlet signal is shown as a volume marker. Scale bars, 1 µm. ***C,*** Distribution of dendritic spine volumes under plasmid lipofection conditions, plotted as a function of binned spine mVenus intensity. Spine volume was estimated by deconvolution (see Methods). mVenus intensity was measured for each spine, normalized by the estimated spine volume, and averaged across spines within each dendrite. The mean mVenus intensity per unit volume at 50 ng/mL lipofection was defined as 10 a.u./μm³. Six lipofection concentrations (2–500 ng/mL) are shown in different colors. No A/C heterodimerizer was applied. Kruskal-Wallis test (*H* = 81, *p* = 5.5 × 10^-16^) followed by Mann-Whitney test against neurons expressing cell-filling mScarlet alone (N (spines/dendrites) = 69/6). -10^0^ a.u./μm^3^, N = 85/10, *U* = 2853, *p* = 0.77; 10^0^-10^0.5^ a.u./μm^3^, N = 53/5, *U* = 1698, *p* = 0.50; 10^0.5^-10^1^ a.u./μm^3^, N = 34/5, *U* = 588, *p* = 4.1 × 10^-5^; 10^1^-10^1.5^ a.u./μm^3^, N = 33/5, *U* = 310, *p* = 3.1 × 10^-9^; 10^1.5^ a.u./μm^3^, N = 17/3, *U* = 96, *p* = 1.0 × 10^-7^. ***D,*** Example confocal images of dendrites expressing mVenus-FKBP-K7GEF via AAV transduction. Sparsely expressed mScarlet, by using a double-floxed inverted open (DIO) reading frame system in conjunction with low Cre expression, is shown as a volume marker. Scale bars, 1 µm. ***E,*** Distribution of dendritic spine volumes under AAV transduction conditions, plotted as a function of binned spine mVenus intensity as in (***C***). Four different AAV concentrations (3.0 × 10^8^ -1.0 × 10^10^ GC/mL) are shown in different colors. Kruskal-Wallis test (*H* = 55, *p* = 2.7 × 10^-11^) followed by Mann-Whitney test against neurons expressing cell-filling mScarlet alone. -10^0^ a.u./μm^3^, N (spines/dendrites) = 60/5, *U* = 1946, *p* = 0.56; 10^0^-10^0.5^ a.u./μm^3^, N = 37/5, *U* = 1256, *p* = 0.89; 10^0.5^-10^1^ a.u./μm^3^, N = 52/7, *U* = 751, *p* = 4.8 × 10^-8^; 10^1^-10^1.5^ a.u./μm^3^, N = 11/2, *U* = 95, *p* = 7.2 × 10^-5^, ***F,*** Representative images of highly branched dendritic spine (yellow arrow heads). Scale bars, 1 µm. ***G,*** Fraction of highly branched spines per dendrite expressing mVenus–FKBP–K7GEF via lipofection (grey) or AAV (red). Statistical analysis was performed using Mann-Whitney test with Bonferroni correction. Lipofection -10^0.5^ a.u./μm^3^ (N (dendrites) = 15) vs Cell-fill only (N = 7), *U* = 51, *p* = 0.91; AAV -10^0.5^ a.u./μm^3^ (N = 10) vs Cell-fill only, *U* = 34, *p* = 0.82; Lipofection 10^0.5^- a.u./μm^3^ (N = 13) vs Cell-fill only, *U* = 0.0, *p* = 3.4 × 10^-4^; AAV 10^0.5^- a.u./μm^3^ (N = 9) vs Cell-fill only, *U* = 2.5, *p* = 1.6 × 10^-3^; -10^0.5^ a.u./μm^3^ lipofection vs AAV, *U* = 60, *p* = 0.65; 10^0.5^- a.u./μm^3^ lipofection vs AAV, *U* = 68, *p* = 0.90. \**p* < 0.05; \*\**p* < 0.01; n.s., not significant.

**Figure 5.**
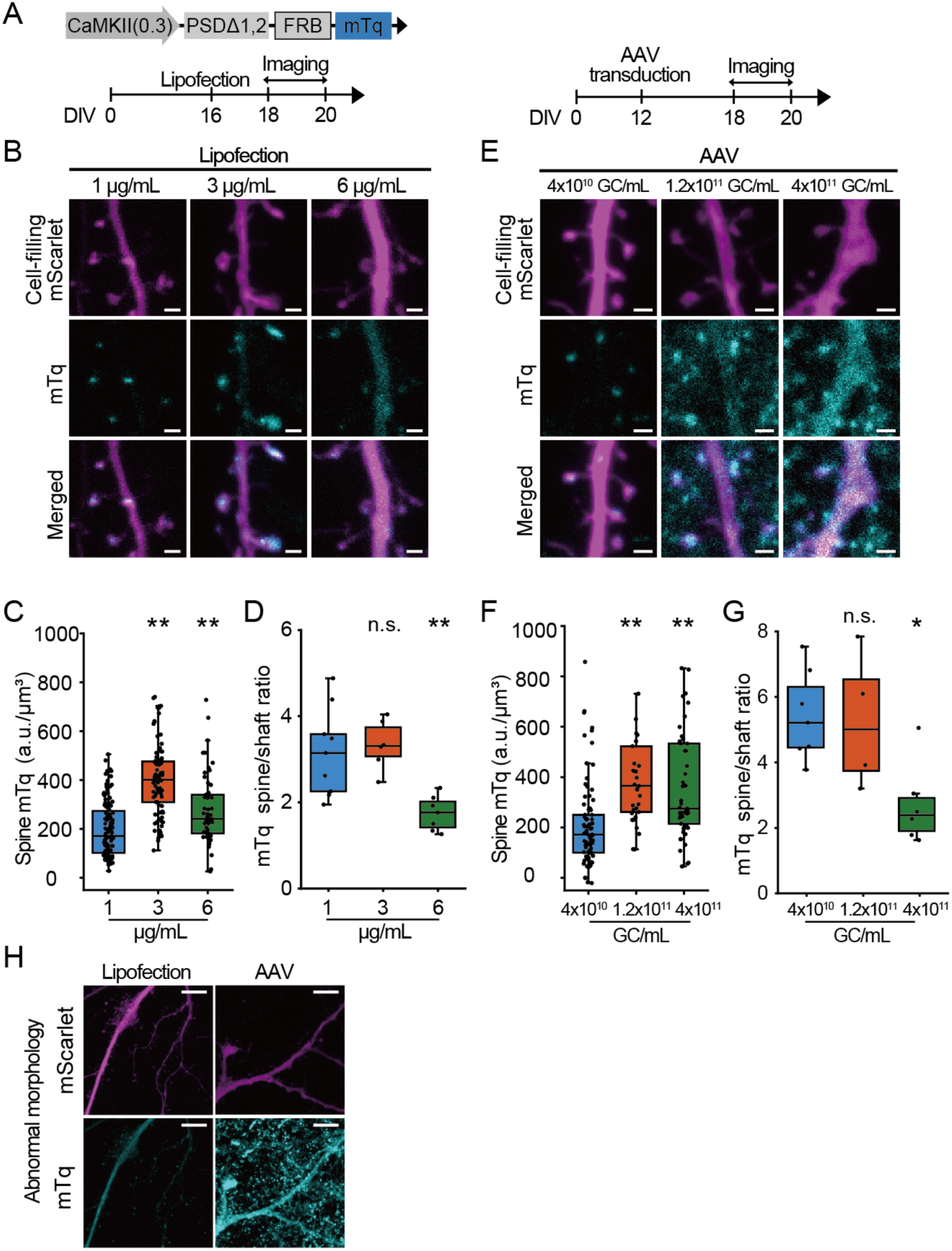
PSDΔ1,2–FRB–mTq induces concentration-dependent spillover into the dendritic shaft and morphological abnormalities. ***A,*** Construct and experimental schedule. Two delivery methods were used: lipofection (***B-D***) and AAV (***E-G***). ***B,*** Example confocal images of dendrites expressing PSDΔ1,2-FRB-mTq via plasmid lipofection. mScarlet signal is shown as a volume marker. Scale bars, 1 µm. ***C,*** Distribution of mTq signal intensity in dendritic spines across lipofection concentrations (1–6 μg/mL). mTq intensity was normalized to the estimated volume of each spine. Each point represents an individual dendritic spine. N (spines/dendrites/neurons) = 111/9/3 (1 μg/mL), 90/6/2 (3 μg/mL), 56/7/5 (6 μg/mL).Statistical comparisons were performed using Kruskal-Wallis test (*H* = 86, *p* = 2.6 × 10^-19^) followed by Mann-Whitney test with Bonferroni correction. 1 vs 3 μg/mL, *U* = 1339, *p* = 4.9 × 10^-19^; 1 vs 6 μg/mL, *U* = 2001, *p* = 1.8 × 10^-4^. ***D,*** Ratio of mTq signal intensity between dendritic spines and shafts within the same dendrite, across lipofection concentrations (1–6 μg/mL). Signals were normalized by their estimated volumes. Each point represents an individual dendrite. Statistical comparisons were performed using the Kruskal-Wallis test (*H* = 12, *p* = 2.6 × 10^-3^) followed by Mann–Whitney test with Bonferroni correction. 1 vs 3 μg/mL, *U* = 23, *p* = 0.69; 1 vs 6 μg/mL, *U* = 59, *p* = 2.1 × 10^-3^. ***E,*** Example confocal images of dendrites expressing PSDΔ1,2-FRB-mTq via AAV transduction. Sparsely expressed mScarlet, by using a double-floxed inverted open (DIO) reading frame system in conjunction with low Cre expression, is shown as a volume marker. Scale bars, 1 µm. ***F,*** Distribution of mTq signal intensity in dendritic spines across AAV concentrations (4.0 × 10^10^ -4.0 × 10^11^ GC/mL), as in (***C***). N (spines/dendrites/neurons) = 83/7/7 (4.0 × 10^10^ GC/mL), 33/4/4 (1.2 × 10^11^ GC/mL), 53/6/6 (4.0 × 10^11^ GC/mL). Kruskal-Wallis test (*H* = 38, *p* = 5.2 × 10^-9^) followed by Mann-Whitney test with Bonferroni correction. 4.0 × 10^10^ vs 1.2 × 10^11^ GC/mL, *U* = 515, *p* = 1.7 × 10^-7^; 4.0 × 10^10^ vs 4.0 × 10^11^ GC/mL, *U* = 1125, *p* = 1.6 × 10^-6^. ***G,*** Ratio of mTq signal intensity between dendritic spines and shafts within the same dendrite, across AAV concentrations, as in (***D***). Kruskal-Wallis test (*H* = 7.9, *p* = 0.019) followed by Mann-Whitney test with Bonferroni correction. 4.0 × 10^10^ vs 1.2 × 10^11^ GC/mL, *U* = 15, *p* = 0.92; 4.0 × 10^10^ vs 4.0 × 10^11^ GC/mL, *U* = 39, *p* = 8.2 × 10^-3^. ***H,*** Representative confocal images of neuronal morphology in neurons transfected with PSDΔ1,2–FRB–mTq by lipofection (6 μg/mL) and infected with AAV-PSDΔ1,2–FRB–mTq (4 × 10¹¹ GC/mL). Scale bars, 10 μm. \**p* < 0.05; \*\**p* < 0.01; n.s., not significant.

Using DNA lipofection at DIV16, we varied the amount of mVenus–FKBP–K7GEF plasmid across six concentrations (2-500 ng/mL), with analyses performed at DIV19 (Fig. 4A, left). The mean mVenus signal intensity per spine volume (see Methods) at 50 ng/mL lipofection was defined as 10 a.u./μm³. When the expression exceeded 10^0.5^ a.u./μm³, the distribution of spine volumes shifted significantly upward even without A/C heterodimerizer, whereas no significant shift was observed below 10^0.5^ a.u./μm³ (Fig. 4B, C). In AAV-based experiments with transduction at DIV11 (Fig. 4A, right), spine volume distributions were likewise shifted at DIV19 only when mVenus–FKBP–K7GEF expression exceeded 10^0.5^ a.u./μm³ (Fig. 4D, E). The comparable expression ranges without unintended effects on spine volume between AAV and lipofection suggest that these effects are primarily determined by expression level rather than by expression duration or delivery method. An increase in highly branched spines, previously reported with Kalirin-7 overexpression (Ma et al., 2008; Penzes et al., 2003), was also observed in both DNA lipofection and AAV, but only when mVenus–FKBP–K7GEF expression exceeded 10^0.5^ a.u./μm³ (Fig. 4F, G). From these results, we considered 10^0.5^ a.u./μm³ as the maximum expression level of mVenus–FKBP–K7GEF without detectable effects on neuronal morphology.

Similarly, PSDΔ1,2–FRB–mTq was expressed by DNA lipofection at three plasmid concentrations (1-6 μg/mL; Fig. 5A, left). Increasing the concentration from 1 to 3 μg/mL significantly elevated PSDΔ1,2–FRB–mTq levels in dendritic spines (the mean spine mTq concentration at 1 μg/mL in lipofection was defined as 200 a.u./μm³) without altering the shaft-to-spine expression ratio (Fig. 5B-D). In contrast, at 6 μg/mL, mTq signal per spine volume did not increase, whereas the spine-to-shaft ratio decreased, suggesting spillover of PSDΔ1,2–FRB from spines to dendritic shafts (Fig.5B-D). Consistent with this, in AAV-transduced neurons (Fig. 5A, right), spine mTq plateaued at ∼400 a.u./μm³ on average, despite increasing expression levels, while the spine-to-shaft ratio decreased, consistent with spillover (Fig. 5E-G). Because SynK induces spine enlargement by recruiting FKBP–K7GEF to PSDΔ1,2-FRB localized in dendritic spines, excessive expression of PSDΔ1,2-FRB in shafts is likely to compromise the efficacy of the tool. Moreover, at expression levels that induced spillover, neurons frequently exhibited abnormal morphologies, including highly-branched spines and lamellipodia like structures (Fig. 5H) and an aberrant increase in small spines and filopodia, suggesting that mislocalization of PSDΔ1,2-FRB adversely affects cellular integrity. Accordingly, we defined 3 μg/mL (lipofection) and 1.2 × 10¹¹ GC/mL (AAV) as the upper limits of the expression range without detectable adverse effects for PSDΔ1,2–FRB–mTq.

### IRESv10-linked SynK induces spine enlargement across a broad expression range without structural abnormalities

To maximize the efficacy of SynK-mediated spine enlargement, each component should be expressed at sufficiently high levels within its safe expression range. Given the substantial difference in expression ranges associated with off-target morphological effects between the two components, we designed a single-construct SynK by linking them with IRES variants (IRESv) (Koh et al., 2013) that allow tuning of their relative expression levels (Fig. 6A-C). Consistent with previous reports, PSDΔ1,2–FRB–mTq expression levels did not differ significantly across IRES variants (Fig. 6D), whereas the PSDΔ1,2–FRB–mTq to mVenus–FKBP–K7GEF expression ratio in dendritic spines, normalized to IRESwt, was 0.32 ± 0.03 for IRESv10 and 0.05 ± 0.03 for IRESv12 (Fig. 6E). This ratio did not differ significantly when the same IRES variant was used at different plasmid concentrations (1 or 3 μg/mL lipofection), indicating that relative expression levels are robustly determined by each IRES variant (Fig. 6E). Together, these results demonstrate that IRES variants enable stable and predictable control of relative protein expression in neurons, consistent with observations in other cell types.

**Figure 6.**
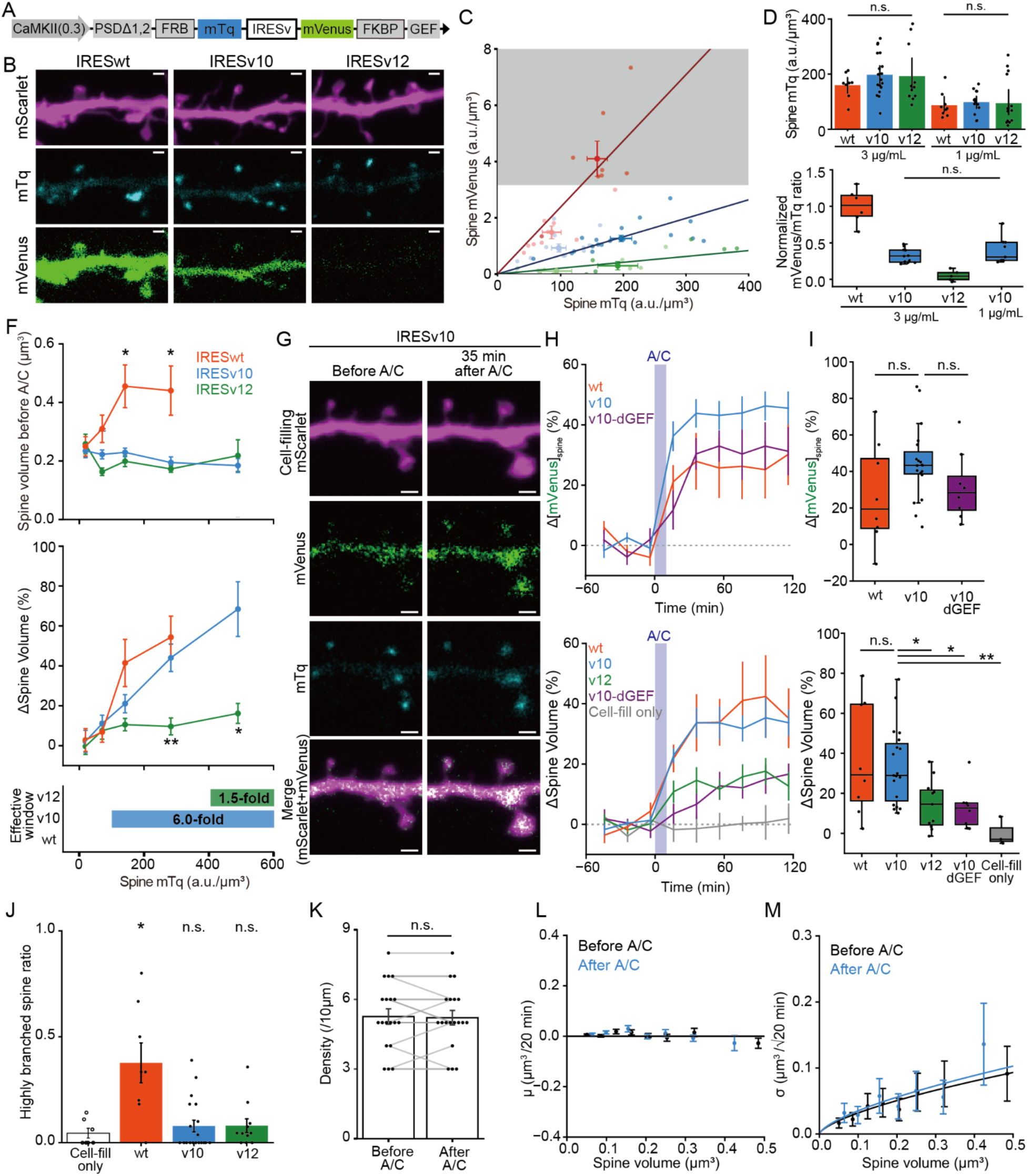
Combining two SynK components using IRESv10 induces dendritic spine enlargement across a broad expression range without detectable structural abnormalities under plasmid lipofection. ***A,*** Schematic of a single-vector construct combining two SynK components via IRES variants (IRESv). Three variants with different translation efficiencies were used: IRESwt, IRESv10 and IRESv12. ***B,*** Representative confocal images of dendrites expressing each IRES construct, delivered via plasmid lipofection: IRESwt (left), IRESv10 (middle) and IRESv12 (right). Scale bars, 1 μm. ***C,*** Distributions of mTq and mVenus intensities in dendritic spines under six conditions (v12, v10, and wt at 1 and 3 μg/mL) before adding A/C heterodimerizer (-5 min). Signals were measured per spine and averaged per dendrite. Gray shading indicates mVenus concentrations that induce unintended baseline changes in spine volume (Fig. 4; 10^0.5^ a.u./μm^3^). Only dendrites with background-subtracted signal intensity ≥ 0 are shown. Mean ± s.e.m. for each condition are indicated by crosshairs. Linear regression lines are shown for each IRES variant. ***D,*** Distributions of mTq intensities in dendritic spines under six conditions (v12, v10, and wt at 1 and 3 μg/mL lipofection). Only dendrites with background-subtracted signal intensity ≥ 0 are shown. N (dendrites/neurons) = 8/6 (wt, 3 μg/mL), 19/11 (v10, 3 μg/mL), 11/5 (v12, 3 μg/mL), 9/7 (wt, 1 μg/mL), 13/9 (v10, 1 μg/mL) and 14/8 (v12, 1 μg/mL). Statistical comparisons were made using the Kruskal-Wallis test (1 μg/mL, *H* = 0.78, *p* = 0.68; 3 μg/mL, *H* = 1.8, *p* = 0.41). ***E,*** Expression ratio of mVenus to mTq for the three IRES variants. Signals were measured per spine and averaged per neuron. Statistical comparison between v10 (3 μg/mL) and v10 (1 μg/mL) was performed using the Mann–Whitney test (*U* = 35, *p* = 0.29). ***F,*** The concentration range at which SynK constructs linked via each IRES variant induce A/C-dependent spine enlargement without affecting baseline spine volume. (Top) Averaged dendritic spine volume before A/C heterodimerizer addition, plotted for each IRES variant across expression levels measured by spine mTq intensity. Data were binned in doubling intervals, with the first bin defined as ≤50 a.u., and the mean ± s.e.m. for each bin is shown. Statistical comparisons were performed using the Mann–Whitney test, comparing each variant to cell-fill–only neurons within each bin, with Bonferroni correction. For wt, - 50 a.u./μm³, N (spines) = 36, U= 1469, *p* = 0.13; 50-100 a.u./μm³, N = 43, *U* = 1853, *p* = 0.027; 100 - 200 a.u./μm³, N = 44, *U* = 2086, *p* = 8.3 × 10^-4^; 200 - 400 a.u./μm³, N = 19, *U* = 954, *p* = 2.5 × 10^-3^. For v10, - 50 a.u./μm³, N = 37, *U* = 1586, *p* = 0.041; 50-100 a.u./μm³, N = 69, *U* = 2775, *p* = 0.094; 100 - 200 a.u./μm³, N = 107, *U* = 4303, *p* = 0.064; 200 - 400 a.u./μm³, N = 51, *U* = 1779, *p* = 0.92; 400 - 600 a.u./μm³, N = 12, *U* = 457, *p* = 0.57. For v12, - 50 a.u./μm³, N = 49, *U* = 1851, *p* = 0.38; 50- 100 a.u./μm³, N = 54, *U* = 1806, *p* = 0.77; 100 - 200 a.u./μm³, N = 79, *U* = 2844, *p* = 0.65; 200 - 400 a.u./μm³, N = 53, *U* = 1905, *p* = 0.70; 400 - 600 a.u./μm³, N = 18, *U* = 646, *p* = 0.80. (Middle) Percentage change in dendritic spine volume between 35 and 75 min following A/C heterodimerizer administration, plotted in the same manner. Statistical comparisons were performed using the Mann–Whitney test, comparing each variant to IRESv10 within each bin, with Bonferroni correction. For wt, - 50 a.u./μm³, *U* = 659, *p* = 0.94; 50-100 a.u./μm³, *U* = 1352, *p* = 0.43; 100 – 200 a.u./μm³, *U* = 2723, *p* = 0.13; 200 - 400 a.u./μm³, *U* = 563, *p* = 0.30. For v12, - 50 a.u./μm³, *U* = 872, *p* = 0.77; 50-100 a.u./μm³, *U* = 1718, *p* = 0.46; 100 - 200 a.u./μm³, *U* = 3927, *p* = 0.41; 200 – 400 a.u./μm³, *U* = 595, *p* = 8.8 × 10^-7^; 400 - 600 a.u./μm³, *U* = 33, *p* = 1.6 × 10^-3^. (Bottom) For each bin, the mean spine volume before A/C and the mean change in spine volume were calculated. For each IRES variant, bins exhibiting ≥15% increase in spine volume without a significant increase in baseline dendritic spine volume were identified, and the corresponding spine mTq concentration range (minimum to maximum) is indicated by markers. ***G,*** Representative confocal image of a dendrite expressing PSDΔ1,2–FRB–mTq–IRESv10–mVenus–FKBP–K7GEF before (−5 min) and after (35 min) application of A/C heterodimerizer. mScarlet signal is shown as a volume marker. Scale bars, 1 µm. ***H, I,*** Averaged time courses (***G***) and summary quantification (***H***) of changes in dendritic spine mVenus concentration in quantified as the mVenus-to–mScarlet ratio (top) and spine volume (bottom) from neurons expressing SynK constructs linked via IRESwt, IRESv10, or IRESv12, as well as an IRESv10 GEF-dead mutant (dGEF), following lipofection at 3 μg/mL. mVenus signals for IRESv12 were not plotted due to unreliable detection. Data in (***G***) are shown as the mean ± s.e.m. Values in (***H***) represent the average between 35 and 115 min after A/C heterodimerizer administration. N (dendrites/neurons) =8/6 (wt), 19/11 (v10), 11/5 (v12), 8/5 (v10 dGEF), and 3/2 (cell-fill only). The median (horizontal line), quartiles (boxes), and range within 1.5 times the interquartile range (whiskers) are denoted. Statistical comparisons were made using the Kruskal-Wallis test (top, *H* = 3.7, *p* = 0.16; bottom, *H* = 16, *p* = 2.5 × 10^-3^) followed by Mann–Whitney test against neurons expressing IRESv10 with Bonferroni correction (bottom, wt, *U* = 76, *p* = 1.0; v12, *U* = 164, *p* = 0.011; v10-dGEF, *U* = 126, *p* = 6.5 × 10^-3^, Cell-fill only, U = 23, *p* = 1.3 × 10^-3^). ***J,*** Ratio of highly branched spines per dendrite under 3 μg/mL plasmid lipofection. Statistical comparisons were performed using the Kruskal–Wallis test (*H* = 11, *p* = 9.7 × 10^-3^), followed by Mann–Whitney tests against neurons expressing cell-filling mScarlet alone (without SynK). IRESwt, *U* = 5, *p* = 8.0 × 10^-3^; IRESv10, *U* = 62, *p* = 0.80; IRESv12, *U* = 31, *p* = 0.51. Data for cell-fill only were reproduced from Fig.4G for comparison. ***K,*** Spine density in neurons expressing SynK linked via IRESv10 following 3 μg/mL lipofection, before and after A/C heterodimerizer administration, quantified per 10 μm of dendrite. N (dendrites/neurons) =19/11 (v10). Wilcoxon signed-rank test (*W* = 12, *p* =0.71). ***L, M,*** Intrinsic spine fluctuation of neurons expressing PSDΔ1,2–FRB–mTq–IRESv10–mVenus–FKBP–K7GEF, before and after A/C heterodimerizer administration, each calculated per 20-min interval. Each plotted point represents the mean (***L***) and the s.d. (***M***) of spine-head volume changes in 21 pooled spines with similar baseline volumes. Error bars represent s.e.m. values (***L***) and the 95% confidence intervals of the estimated s.d. (***M***). Data are fitted by a zero (***L***) and to the 2/3 power of the baseline spine volume (***M***). \**p* < 0.05; \*\**p* < 0.01; n.s., not significant.

We next analyzed structural abnormalities before A/C administration and A/C-induced spine enlargement in SynK constructs linked by each IRES variant, while varying the DNA lipofection concentration within a range (1–3 μg/mL) in which PSDΔ1,2–FRB–mTq does not cause morphological abnormalities (Fig. 5). Before A/C heterodimerizer administration, IRESwt, under which mVenus–FKBP–K7GEF expression more readily exceeded the safe range (Fig. 6C) and exhibited increased spine volume (Fig. 6F, top). In contrast, within a similar DNA concentration range, neither mVenus–FKBP–K7GEF expression exceeding this range nor these unintended effects were observed with IRESv10 or IRESv12 (Fig. 6C and F, top).

Following A/C heterodimerizer administration, mVenus–FKBP–K7GEF translocation to dendritic spines was comparably observed in IRESwt and IRESv10, whereas in IRESv12 the mVenus signal was largely undetectable, precluding reliable assessment of translocation (Fig. 6G-I). Consistently, spine volume increases were comparable between IRESwt and IRESv10, whereas IRESv12 showed a significantly smaller effect, likely reflecting insufficient mVenus–FKBP–K7GEF expression (Fig. 6F, middle and G-I).

Across expression titrations for each variant, we defined the effective concentration window (dynamic range) as the fold range between the highest and lowest concentrations that showed no detectable alterations in spine volume prior to A/C heterodimerizer administration while still inducing >15% spine enlargement in an A/C-dependent manner. This dynamic range was largest for IRESv10 (6.0-fold); no such range was observed for IRESwt, and only a narrow range was observed for IRESv12 (1.5-fold). Thus, IRESv10 provides the most balanced expression of the two components, achieving optimal efficacy with minimal baseline morphological perturbation among the variants tested.

In the IRESv10 construct, replacing the GEF domain with a catalytically inactive mutant (dGEF) (Penzes et al., 2001) did not affect A/C heterodimerizer–dependent accumulation in spines (Fig. 6H and I, top panels), but significantly reduced the extent of spine enlargement (Fig. 6H and I, bottom panels), suggesting that spine enlargement is dependent on GEF activity. In IRESv10, the proportion of highly branched spines was not significantly increased compared to control neurons lacking SynK expression, whereas it was elevated in IRESwt (Fig. 6J). We also examined spine density or intrinsic spine fluctuation before and after A/C heterodimerizer administration, finding no significant unintended changes (Fig. 6K-M).

### AAV-based validation confirms that IRESv10-linked uniSynK reliably induces spine enlargement with minimal unintended morphological abnormalities

Finally, we termed the IRESv10-based construct uniSynK and compared it with the original split two-component SynK using an AAV-based approach (Fig. 7A). To visualize neuronal morphology, mScarlet was sparsely expressed using low levels of Cre recombinase, whereas split SynK or uniSynK was delivered independently and broadly expressed across infected neurons. When the two components of the original split SynK system were delivered at a 50:1 ratio, the resulting expression ratio between PSDΔ1,2–FRB-mTq and mVenus-FKBP-K7GEF was, on average, comparable to that achieved with IRESv10 in lipofection condition (Fig. 6E), but exhibited substantial variability (Fig. 7B, left and C). To further examine this variability, cells were categorized into three groups (Groups 1–3) based on their expression ratios (Fig. 7C), with Group 2 corresponding to cells within the IRESv10-equivalent range (Fig. 6E; 0.21–0.48).

**Figure 7.**
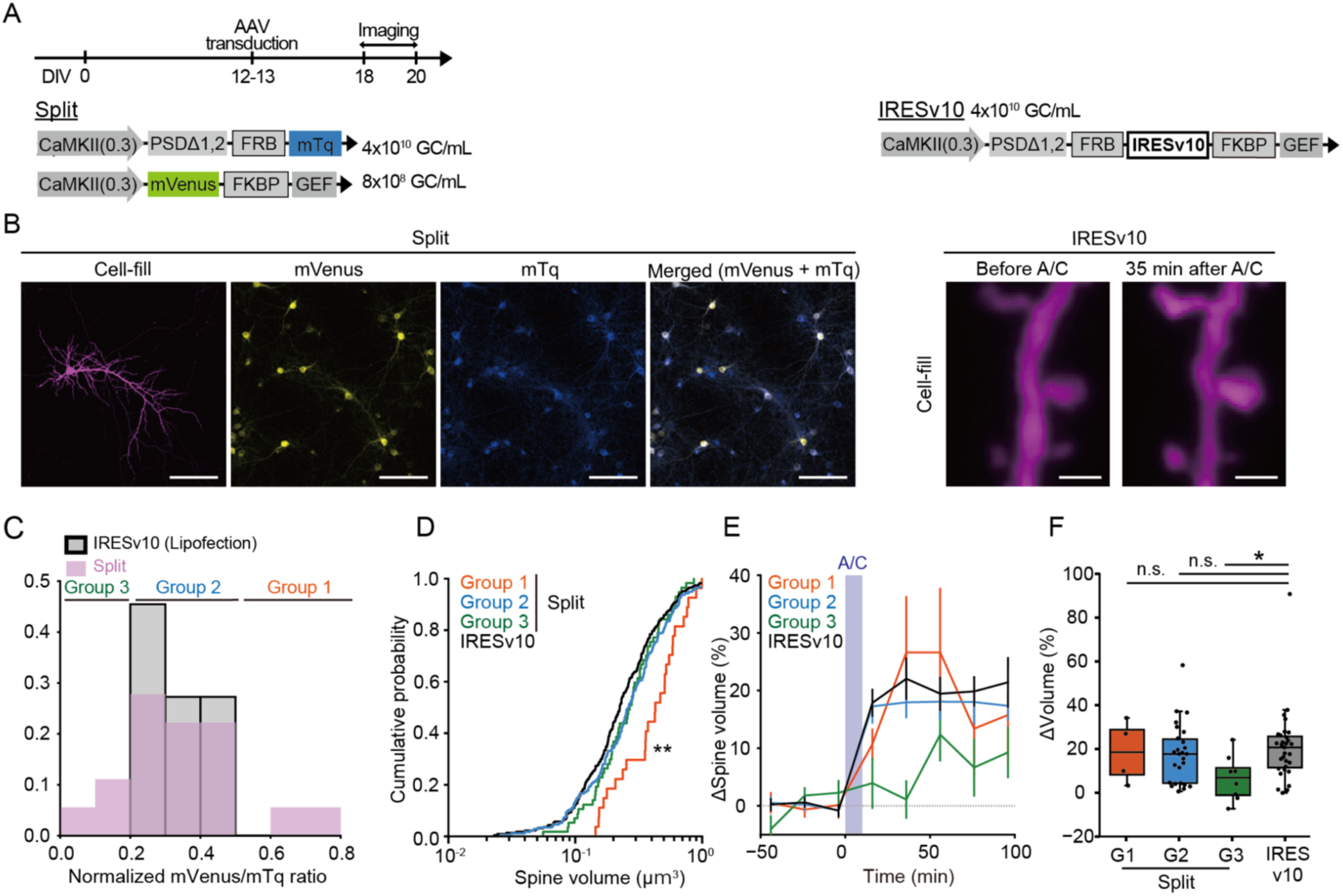
AAV-based validation confirms that IRESv10-linked uniSynK reliably induces spine enlargement with minimal unintended morphological abnormalities under baseline conditions. ***A,*** Schematic of uniSynK (single construct linked via IRESv10) and split SynK (two constructs). Constructs were delivered using AAV. ***B,*** (Left) Representative confocal images of dendrites expressing split SynK. For visualization purpose only, higher concentrations were used while maintaining the same delivery ratio (PSDΔ1,2-FRB-mTq, 1 × 10¹¹ GC/mL; mVenus-FKBP-K7GEF, 2 × 10⁹ GC/mL). Scale bars, 50 µm. (Right) Representative confocal images of dendrites expressing uniSynK before (−5 min) and after (35 min) application of A/C heterodimerizer. mScarlet signal is shown as a volume marker. Scale bars, 1 µm. ***C,*** Distribution of the mVenus-to-mTq expression ratio in spines for the split construct. Ratios were calculated for individual spines and averaged per cell for plotting. Cells were classified into three groups based on these ratios: within the IRESv10 range defined under 3 μg/mL plasmid lipofection (including both mTq and mVenus; Fig. 6E; group 2), above this range (group 1), and below this range (group 3). ***D,*** Cumulative distributions of dendritic spine volumes under four conditions: uniSynK and split SynK groups 1–3. N (spines/dendrites/neurons) = 227/29/16 (IRESv10), 27/4/2 (group 1), 175/26/12 (group 2) and 57/8/4 (group 3). Statistical comparisons were performed using the Kolmogorov–Smirnov (K–S) test against the uniSynK condition with Bonferroni correction.Group 1, *D* = 0.44, *p* = 9.6 × 10^-5^; Group2, *D* = 0.12, *p* = 0.097; Group3, *D* = 0.13, *p* = 0.39. ***E, F,*** Averaged time courses (***E***) and summary quantification (***F***) of changes in spine volume from neurons expressing uniSynK and split SynK groups 1–3. Data in (***E***) are shown as the mean ± s.e.m. Values in (***F***) represent the average between 15 and 95 min after A/C heterodimerizer administration. N (dendrites/neurons) = 29/16 (uniSynK, IRESv10), 4/2 (split SynK group1), 26/12 (split SynK group2) and 8/4 (split SynK group 3). The median (horizontal line), quartiles (boxes), and range within 1.5 times the interquartile range (whiskers) are denoted. Statistical comparisons were made using Mann–Whitney test against the uniSynK condition with Bonferroni correction. Group 1, *U* = 61, *p* = 0.98; Group 2, *U* = 443, *p* = 0.52; Group 3, *U* = 198, *p* = 8.6 × 10^-3^. \**p* < 0.05; \*\**p* < 0.01; n.s., not significant.

In Group 1, where mVenus-FKBP-K7GEF expression was relatively high, increased spine volume was evident prior to A/C application (Fig. 7D), consistent with the IRESwt condition (Fig. 6F), suggesting baseline overactivation. In contrast, in Group 3, where mVenus–FKBP–K7GEF expression was relatively low, no significant changes in baseline spine volume were observed (Fig. 7D), but A/C-induced spine enlargement was attenuated (Fig. 7E, F), consistent with the IRESv12 condition (Fig. 6G, H). In Group 2, significant spine enlargement was induced without detectable unintended effects in baseline conditions (Fig. 7D-F). Together, these results further confirm that precise control of the expression ratio between the two components is critical for achieving safe and effective induction of spine enlargement.

In contrast, a single AAV construct linking the two components via IRESv10 (uniSynK; 4.6 kb)—lacking mVenus and mTq due to packaging constraints—produced significantly fewer morphological abnormalities under baseline conditions than split Group 1, and levels comparable to Groups 2 and 3 (Fig. 7D). Moreover, uniSynK exhibited A/C-dependent spine enlargement comparable to Groups 1 and 2, and greater than Group 3 (Fig. 7E, F), demonstrating stable and safe induction of spine enlargement. Across all dendrites included in the analysis, 71% of dendrites expressing uniSynK exhibited no significant deviation from baseline levels prior to A/C addition (defined as within ±2 SD of the control distribution) while still showing a mean A/C-dependent spine volume increase of >10%, compared with 41% in the split SynK condition. These results indicate that the combined vector improves the reliability of inducible spine enlargement while minimizing unintended baseline morphological abnormalities, likely by constraining variability in the relative expression levels of the two components.

## Discussion

In this work, we present uniSynK, a new tool to induce dendritic spine enlargement by integrating the two separate components of the original SynK (Sawada et al., 2024) into a single construct. When multiple AAVs are used, it is inherently challenging to achieve coordinated expression of all components within their respective optimal ranges across cells (Chen et al., 2013; Grieger et al., 2006; Maturana et al., 2023; Tian et al., 2009). In the original SynK system, this limitation was particularly pronounced for the FKBP–K7GEF component, which required low expression levels to function properly; accordingly, its AAV titer had to be reduced, making it difficult to achieve sufficient transduction across a large population of neurons.

To address this, we first quantitatively defined the expression-level thresholds at which each component produced unintended effects (Fig. 4 and 5). We then incorporated IRES variants with graded translational efficiencies (Koh et al., 2013) to fine-tune the expression ratio of the two components, enabling stable and robust expression within the optimal functional range (Fig. 6). Notably, we found that these IRES variants provided stable relative expression levels in neurons as well (Fig. 6E). The resulting IRESv10-based construct, uniSynK, enabled A/C-dependent spine enlargement over a relatively broad concentration range without evident off-target effects (Fig. 6E and 7).

Precise control of relative protein expression is a central requirement when designing co-expression systems. One conventional approach is to combine promoters of different strengths; however, placing multiple promoters in close proximity within a single vector can cause transcriptional interference and thereby compromise the predictability of transgene expression (Eszterhas et al., 2002; Schlatter et al., 2005). Splicing-based strategies can also be used to modulate expression ratios between two genes (Aebischer-Gumy et al., 2024; Fallot et al., 2009). However, splice-site usage depends on both local sequence context, including cis-acting splicing enhancers and silencers, and the cellular repertoire of splicing regulatory factors, limiting the general applicability of this approach for maintaining stable expression ratios across different transgene pairs (Renaud-Gabardos et al., 2015).

2A peptides and IRESs are widely used alternatives for co-expressing multiple genes from a single mRNA. Because of their short length, 2A peptides are particularly useful under AAV packaging constraints (de Felipe et al., 2006; Ryan et al., 1991), and their utility has also been demonstrated in neurons (Furler et al., 2001). However, incomplete cleavage can generate uncleaved fusion products (Chan et al., 2011; de Felipe et al., 2010; Ho et al., 2013). In the SynK system, such products could potentially cause premature localization of the Kalirin-7 GEF domain to dendritic spines before A/C heterodimerizer addition, increasing the risk of unintended effects. In addition, 2A peptides do not readily enable precise tuning of the relative expression ratio, which becomes problematic when the two components require substantially different expression levels.

We therefore adopted an IRES-based strategy (Renaud-Gabardos et al., 2015), in which the expression ratio can be tuned by selecting IRES variants with different translational efficiencies (Koh et al., 2013). Although the relatively large size of IRES sequences is a potential drawback, we offset this constraint by successfully shortening the construct. Moreover, although the use of IRES variants for quantitative control of relative protein expression in neurons has not been well established, our results demonstrate that IRES-dependent tuning can be achieved in neurons. These findings support IRES variants as a flexible platform for optimizing multi-protein expression systems, particularly when precise ratiometric control is required.

While quantitatively assessing the side effects of PSDΔ1,2-FRB and FKBP-K7GEF expression, we found that each component displayed distinct modes of unintended effects. PSDΔ1,2 has long been used as a postsynaptic marker (Arnold and Clapham, 1999; Fujimoto et al., 2023; Hayashi-Takagi et al., 2015; Sturgill et al., 2009), yet the consequences of its overexpression had not been systematically quantified. We observed that the spine/shaft ratio was maintained within a certain expression range (Fig. 5B-G), within which overall cellular architecture remained largely unaffected. However, once PSDΔ1,2 expression exceeded its postsynaptic localization capacity and ectopically leaked into dendritic shafts, catastrophic abnormalities emerged, highly-branched spines and lamellipodia like structures and an aberrant increase in small spines and filopodia (Fig. 5H). In contrast, Kalirin-7 produced more modulatory effects that scaled in a relatively linear fashion with concentration, increasing average spine volume and promoting branched spines and spinules (Fig. 4). Altered expression levels of Kalirin-7 and PSD proteins have been reported in depression, autism and schizophrenia (Coley and Gao, 2018; Duric et al., 2013; Feyder et al., 2010; Karolewicz et al., 2009; Remmers et al., 2014; Rodríguez-Palmero et al., 2021; Russell et al., 2014), and our quantitative investigation on the graded mode of action may offer new insight into how genetic abnormalities translate into synaptic and, ultimately, behavioral phenotypes.

Although the uniSynK system demonstrated improved performance in cultured neurons, several limitations and opportunities for further refinement remain. First, due to the packaging constraints, it was not possible to include an additional fluorescent reporter in the final AAV-uniSynK construct (Fig. 7), and thus direct visualization of expression efficiency at the population level was not feasible in the present study. However, the IRES-based architecture ensures stable and reproducible expression ratios between the two components (Fig. 6E), which may compensate for this limitation by reducing the need for post hoc selection of cells with appropriate expression levels. Second, while uniSynK was robustly validated in cultured neurons using both lipofection and AAV transduction, its efficacy and safety have not yet been assessed in vivo, where AAV transduction patterns and promoter activity may differ from those observed in vitro. Future in vivo validation will therefore be an important step toward establishing its broader applicability. Third, although integrating the two components into a single construct substantially improves controllability compared to split systems, excessive expression levels can still give rise to unintended effects. Further refinements may reduce such leak effects, for example by introducing additional structural switching mechanisms into Kalirin-7 itself or by enhancing dimerization-dependent translocation efficiency to permit lower expression levels.

Despite these remaining limitations, the conceptual framework established here—quantitative definition of expression thresholds and ratiometric control of multi-component systems—addresses a challenge that extends well beyond the SynK system itself. To date, numerous two-component systems have been developed, ranging from chemically induced translocation (CIT) tools for diverse cell types (Bisaria et al., 2020; Inoue et al., 2005) to neuron-targeted tools for tagging and manipulation (Hyun et al., 2022; Idevall-Hagren et al., 2012; Kim et al., 2020; Lee et al., 2017). However, their use is often limited by the need to fine-tune the expression levels of each component. Here, we provide a practical solution by leveraging IRES variants to precisely control expression ratios, which could be broadly applied to such systems and facilitate their adoption.

In addition to its broader methodological implications, uniSynK provides a practical means to address long-standing biological questions regarding the causal roles of synaptic plasticity. Synaptic alterations have been implicated in a wide range of conditions, including sleep regulation, psychiatric disorders, and brain tumors. Directly testing the functional consequences of spine and synaptic modifications requires tools that perturb synapses themselves, thereby making synapses causal experimental variables. Compared with optogenetic approaches (Goto et al., 2021; Hayashi-Takagi et al., 2015), synapse-targeted chemogenetic manipulation may offer distinct advantages. Structural manipulation of synapses using light can require extended illumination and high power, increasing the risk of phototoxic effects, whereas ligand-dependent systems enable sustained modulation following a single administration without continuous intervention. Moreover, systemic delivery of an otherwise inert chemical dimerizer allows effective manipulation across large brain regions—or even the entire brain—without the need for invasive optical access. When combined with cell type–specific enhancers (Ben-Simon et al., 2025; Challis et al., 2022) and synapse-tagging strategies (Gobbo et al., 2017; Hayashi-Takagi et al., 2015) that selectively label defined synaptic populations, uniSynK could be further extended to manipulate specific synapse subsets, providing a versatile framework for dissecting synapse-specific functions across intact neural circuits.

In summary, uniSynK establishes a generalizable strategy for precise and balanced control of multi-component systems, enabling robust synaptic actuation.

## Acknowledgements

We thank S. Fujii, C. Fujinami, and M. Hama for technical assistance. During manuscript preparation, the authors used OpenAI’s GPT-5 language model for English editing and phrasing refinement under full author supervision. This work was supported by FOREST (JPMJFR231T to T.S.) and CREST (JPMJCR21E2 to H.K.) from JST, Brain/MINDS 2 (JP24wm0625101 to H.K., S.Z. and T.S.) from AMED, KAKENHI (JP23K14385 to T.S.; JP20H05685 to H.K., S.Z. and T.S.) from JSPS, and the World Premier International Research Center Initiative (WPI) from MEXT.

## Author contributions

K.H., H.O., and T.S. designed research; K.H., M.O., S.Z., and T.A. performed research; K.H. and T.S. analyzed data; H.K. provided critical input; K.H. and T.S. wrote the paper; all authors edited it.

